# Specificity Protein 1 is essential for the limb trajectory of ephrin-mediated spinal motor axons

**DOI:** 10.1101/2025.01.07.631735

**Authors:** Pinwen Liao, Ming-Yuan Chang, Wen-Bin Yang, Keefer Lin, Yi-Chao Li, Jian-Ying Chuang, Yi-Hsin Wu, Artur Kania, Wen-Chang Chang, Tsung-I Hsu, Tzu-Jen Kao

## Abstract

The precise organization of neural circuits requires highly specific axon guidance, facilitated by cell-surface guidance receptors on axonal growth cones that help neurons reach their target destinations. Despite a limited repertoire of known guidance receptors and ligands, neural systems achieve complex axonal networks, suggesting that additional regulatory mechanisms exist. One proposed strategy is the co-expression of ligands and their receptors on the same axons, allowing modulation of receptor responsiveness to guidance cues. To investigate this mechanism, we studied the spinal lateral motor column (LMC) motor neurons, which make a binary axon pathfinding decision toward limb targets. We hypothesized that specificity protein 1 (Sp1), a transcription factor, regulates ephrin expression in LMC neurons, thereby modulating receptor functions via *cis*-attenuation to ensure accurate axonal pathfinding. Our results show that Sp1 is indeed expressed in LMC neurons during critical axonal extension periods. Manipulating Sp1 activity disrupted LMC axon trajectory selection, and RNA-Seq analysis indicated that Sp1 regulates genes associated with axon guidance, including ephrins. We found that Sp1 knockdown affected ephrin/Eph *cis*-binding and *trans*-signaling, highlighting Sp1’s role in controlling axonal projections through ephrin gene regulation. Additionally, coactivators p300 and CBP are essential for Sp1’s regulatory function. These findings identify Sp1 as a key transcription factor in LMC neurons, essential for ephrin expression and ephrin/Eph-mediated axon guidance, providing insights into the molecular mechanisms of neural circuit formation.

## Introduction

The complex organization of neural circuits, which depends on highly specific axon guidance, is essential for forming a functional nervous system. Neuronal axonal growth cones need precise guidance to reach their correct targets, a process facilitated by the activation of cell-surface guidance receptors located on the axons. Given the relatively limited number of known protein families, especially those encoding axon guidance receptors and their ligands, questions arise about how the diversity of guidance signals is amplified to direct the intricate axonal networks within most neural circuits. One proposed solution to this challenge is the co-expression of ligands and their receptors on the same axons, a strategy that appears effective in modulating the responsiveness of growth cone receptors to specific guidance cues. However, our understanding of the molecular mechanisms underlying these processes is still limited, largely due to the complexity of most nervous systems under study.

To address the challenges posed by nervous system complexity, we examined a straightforward binary axon pathfinding decision made by spinal lateral motor column (LMC) motor axons as they extend toward the base of the limb. LMC neurons are divided into medial and lateral populations, which grow toward the limb base and then diverge to form ventral and dorsal limb nerves, respectively (Landmesser, 1978). Multiple ligand and receptor families, including ephrins/Ephs, Netrin-1/Dcc (Unc5c), and other supporting systems, have been implicated in guiding LMC axon growth into the limbs, enabling us to explore the mechanism by which receptor functions are finely regulated through receptor/ligand co-expression (Kania and Klein, 2016). Early studies have suggested that ephrins can reduce the activity of co-expressed Eph receptors (known as *cis*-attenuation), limiting the number of Eph receptors available for binding to ligands expressed on other cells (*trans*-binding), thus enhancing the accuracy of axon trajectories (Hornberger et al., 1999; Rashid et al., 2005; Carvalho et al., 2006; Kao and Kania, 2011). Despite these findings, the regulatory machinery underlying *cis*-attenuation remains unclear.

In the lateral motor column (LMC), the expression of Eph receptors is regulated by LIM homeodomain transcription factors Lim1 and Isl1 (Kania and Jessell, 2003; Luria et al., 2008). However, the regulatory mechanisms for co-expressed ephrin in LMC neurons remain unknown. Recently, we observed the expression of specificity protein 1 (Sp1) in the LMC during the period when spinal motor axons grow into the limbs, suggesting that Sp1 may act as a regulator for ephrin gene expression. Sp1, a member of the Sp/KLF transcription factor family, is widely expressed across various mammalian cell types (Briggs et al., 1986) and regulates genes associated with numerous cellular processes (O’Conner et al., 2016). Sp1 activates transcription by recruiting basal transcription machinery and interacting with TFIID complex members, which include the TATA binding protein (TBP) and TBP-associated factors (TAFs) (Hoey et al., 1993; Emili et al., 1994; Saluja et al., 1998). Additionally, Sp1 can associate with epigenetic modifiers such as histone deacetylase 1 (HDAC1) and DNA methyltransferase 1 (DNMT1) to suppress gene expression (Doetzlhofer et al., 1999; Song et al., 2001; Lagger et al., 2003; He et al., 2005; Li et al., 2010). Sp1 knockout in mice results in early embryonic lethality (around day e10.5) with multiple developmental abnormalities, indicating its critical role in development (Marin et al., 1997). Despite this, knowledge of Sp1’s role in central nervous system (CNS) development is still limited. Studies suggest that Sp1 enhances the expression of the NR1 gene, which encodes a key subunit of the N-methyl-D-aspartate receptor, essential for neuronal differentiation (Liu et al., 2004). Furthermore, members of the Sp family (Sp1, Sp3, and Sp4) are thought to collaborate to activate the transcription of cyclin-dependent kinase 5/p35, a protein necessary for brain function (Ross et al., 2002). Sp1 has also been implicated in regulating the expression of the axon guidance ligand Slit2 and modulating Wnt signaling via β-catenin (Mir et al., 2018). Recent studies have shown that astrocyte-expressed Sp1 supports neurite outgrowth and synaptogenesis (Hung et al., 2020), though the role of neuron-specific Sp1 in neural circuit formation remains unclear. Based on this information, we **hypothesize** that Sp1 regulation of ephrin expression in LMC neurons modulates guidance receptor functions through *cis*-attenuation, which is necessary for accurate axonal growth into the limbs.

In this study, we first demonstrate that Sp1 is expressed in LMC neurons during the period of motor axon extension into the limb. Altering Sp1 activity—either by loss or gain of function—disrupts the selection of LMC axon trajectories in opposite ways. Interestingly, our RNA-Seq analysis revealed that Sp1 likely regulates genes associated with axon guidance, including ephrins, suggesting its involvement in controlling spinal motor axon projections through the regulation of ephrin genes. We further show that Sp1 knockdown reduces ephrin/Eph *cis*-binding and diminishes their trans-signaling without affecting Netrin-1, Sema3F, or GDNF *in vitro*. Additionally, Sp1’s coactivators, p300 and CBP, are both necessary for its role in regulating ephrin expression in this context. Together, these findings indicate that Sp1 is a key transcription factor that regulates ephrin gene expression in LMC neurons and subsequently modulates ephrin/Eph *cis*-attenuation, allowing precise control of ephrin/Eph *trans*-signaling and LMC axon trajectory selection.

## Results

### Sp1 expression in LMC motor neurons

The transcription factor Sp1 has been implicated in several cellular processes, including neuronal differentiation, neurite outgrowth, and synaptogenesis (Liu et al., 2004; Hung et al., 2020). To explore the function of Sp1 in motor axon guidance *in vivo* and define its role, we first assessed Sp1 expression in the lateral motor column (LMC) at the time when motor axons invade the limb mesenchyme. This occurs between HH stage 25 and 26 in chick embryos and at e11.5 in mouse embryos (Hamburger and Hamilton, 1951; Tosney and Landmesser, 1985; Kania et al., 2000). We examined Sp1 expression in the lumbar spinal cord of HH stage 25/26 chick embryos and in both brachial and lumbar spinal cords of e11.5 mouse embryos. The dorsal and ventral limb-innervating subpopulations of LMC neurons were identified using the medial LMC marker *Isl1* and the lateral LMC marker *Lim1* (Fig. 1A, B, F, G, K, L). We observed strong expression of *Sp1* mRNA in both medial and lateral LMC neurons in chick and mouse embryos (Fig. 1C, H, M). Additionally, Sp1 coactivators *p300* and *CBP* were also expressed in LMC neurons (Fig. 1D, E, I, J, N, O). No significant differences in *p300* and *CBP* expression were noted between the medial and lateral LMC or between brachial and lumbar spinal cords in chick and mouse embryos.

**Figure 1.**
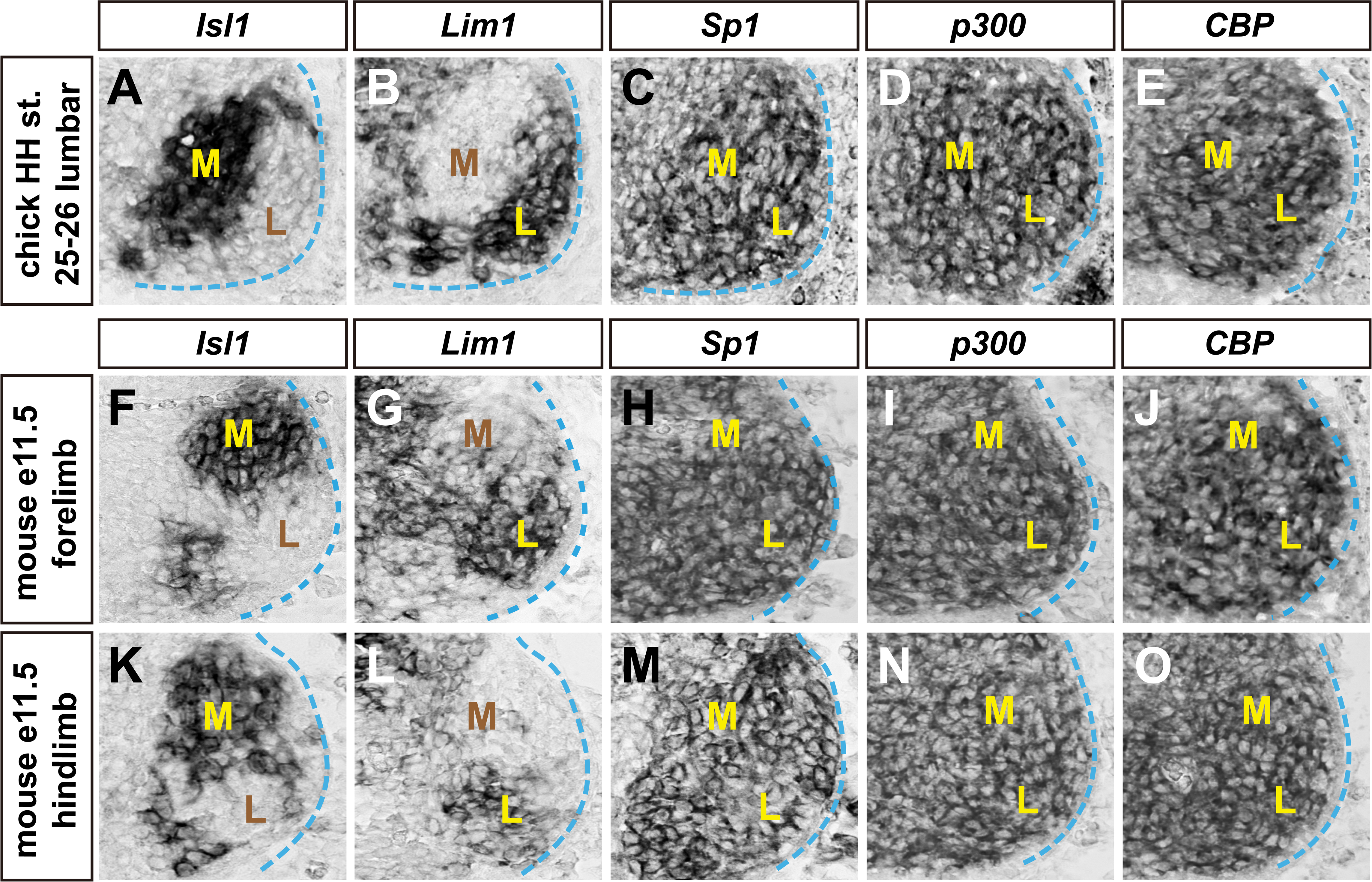
Expression of *Sp1* and its coactivators in chick and mouse LMC motor neurons. **(A-O)** mRNA detection in consecutive sections of spinal cords: all chick sections represent the HH stage 25/26 lumbar spinal cord, while all mouse sections represent the e11.5 lumbar spinal cord. **(A, B)** Detection of ***Isl1*** (A) and ***Lim1*** (B) mRNA in the chick spinal cord, marking the medial and lateral LMC neurons, respectively. **(C-E)** mRNA detection of ***Sp1*** (C) and its coactivators ***p300*** (D) and ***CBP*** (E) in both medial and lateral LMC neurons of the chick. **(F, G, K, L)** mRNA detection of ***Isl1*** (F, K) and ***Lim1*** (G, L) in mouse LMC neurons. **(H-J, M-O)** mRNA detection of ***Sp1*** (H, M), ***p300*** (I, N), and ***CBP*** (J, O) in both medial and lateral LMC neurons in the mouse. M, medial; L, lateral. Scale bar, μm (A-O) 30 μm

### Sp1 is required for limb trajectory selection by LMC axons

To investigate whether Sp1 is involved in determining whether LMC axons select dorsal or ventral limb nerves, we knocked down Sp1 expression by introducing an inhibitory siRNA targeting *Sp1* mRNA (*[Sp1]siRNA*) into LMC neurons. We achieved this by co-electroporating the siRNA along with a *GFP* expression plasmid into the chick lumbar neural tube before LMC neuron specification and before axons entered the limb at HH stage 18/19. We then analyzed GFP^+^ motor axons in the dorsal and ventral nerve branches emerging from the crural plexus at HH stage 28/29 (Kania and Jessell, 2003). The co-electroporation of *[Sp1]siRNA* and *GFP* plasmid resulted in a significant reduction in *Sp1* mRNA and protein levels, compared to embryos that were electroporated with a control *GFP* plasmid or *scrambled [Sp1]siRNA* (Fig. 2A-G). However, this reduction did not lead to noticeable changes in axon outgrowth (Fig. 2K) or in the number of LMC neurons expressing the Foxp1 marker. Additionally, there was no alteration in the proportions of lateral LMC (Foxp1^+^, Isl1^-^) versus medial LMC (Foxp1^+^, Isl1^+^) neurons when compared to embryos treated with the control *GFP* plasmid or scrambled *[Sp1]siRNA* (Fig. 2L, M). The number of electroporated neurons was also similar between the LMC divisions in both the *[Sp1]siRNA* and *GFP* co-expressing groups and the controls (Fig. 2N).

**Figure 2.**
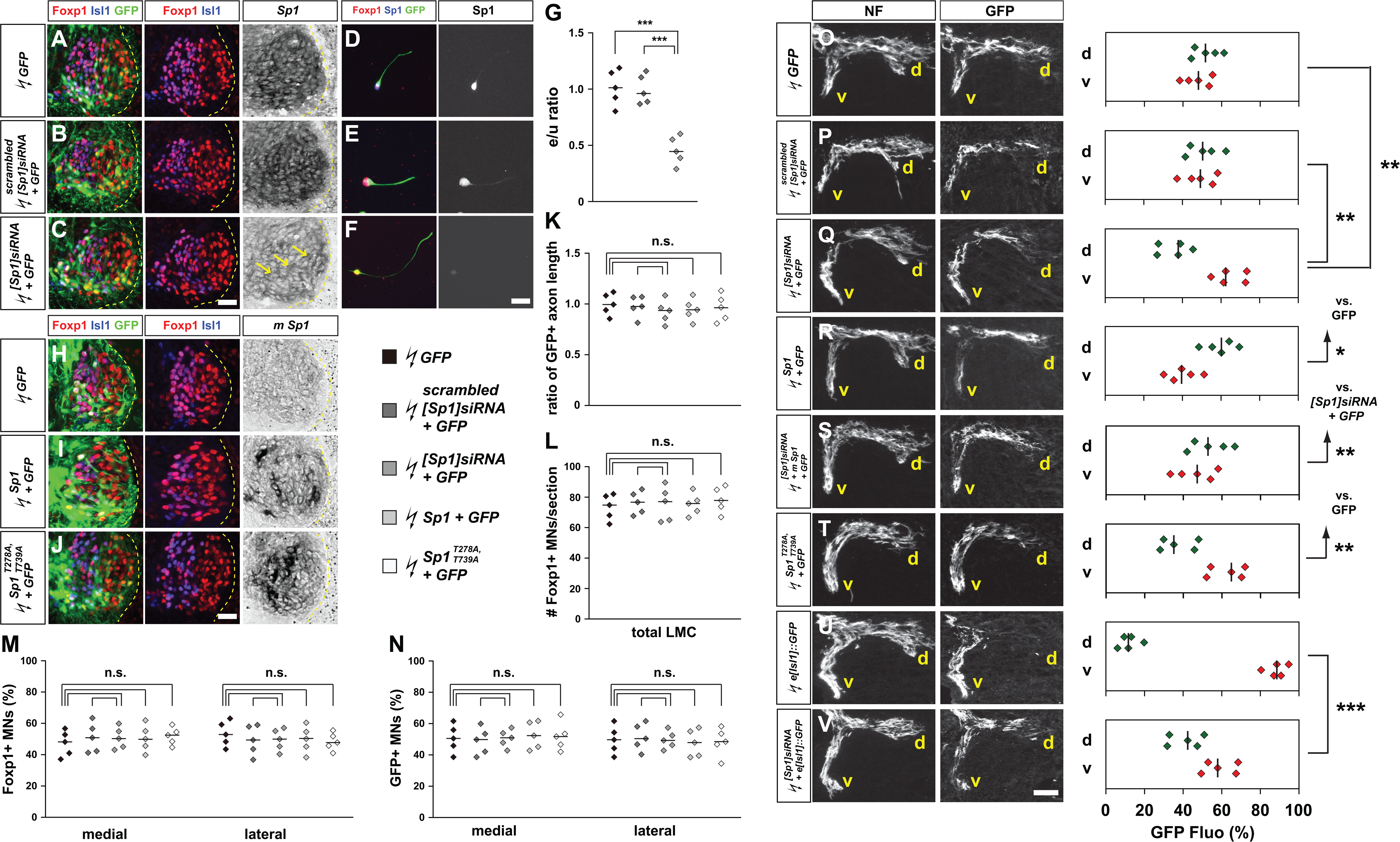
Sp1 function is required for the selection of limb axon trajectory. All images are from chick HH st. 28/29 lumbar levels. (A-C) Detection of Foxp1, Isl1, GFP, and Sp1 in LMC neurons of chick embryos electroporated with either *GFP* (A), *scrambled [Sp1]siRNA* and *GFP* (B), or *[Sp1]siRNA* and *GFP* (C). (D-F) Detection of Foxp1, Sp1, and GFP in cultured LMC neurons from chick embryos electroporated with *GFP* (A), *scrambled [Sp1]siRNA* and *GFP* (B), or *[Sp1]siRNA* and *GFP* (C). (G) Quantification of the effects of *GFP*, *scrambled [Sp1]siRNA* and *GFP*, or *[Sp1]siRNA* + *GFP* electroporation on *Sp1* mRNA levels. The ratio of immunoreactivity signal between the LMC regions of electroporated and un-electroporated contralateral sides (e/u ratio) was measured in at least 15 sections from 5 embryos. (H-J) Detection of Foxp1, Isl1, GFP protein, and mouse *Sp1* mRNA in LMC neurons of chick embryos at HH stage 28/29 electroporated with *GFP* (H), *Sp1* and *GFP* (I), or *Sp1^T278A, T739A^* and *GFP* (J) expression plasmids. (K) Quantification of axon outgrowth effects following electroporation of *GFP*, *scrambled [Sp1]siRNA* and *GFP*, *[Sp1]siRNA* and *GFP*, *Sp1* and *GFP*, or *Sp1^T278A, T739A^* and *GFP*. GFP^+^ LMC axon lengths were compared to the GFP control group, with measurements taken from at least 15 sections of 5 embryos. (L) The number of LMC motor neurons was expressed as the average count of Foxp1**^+^** LMC neurons per section (**#** Foxp1^+^ MNs/section), with 5 embryos per group. (M, N) The number of medial (Foxp1**^+^** Isl1**^+^**) and lateral (Foxp1**^+^** Isl1**^-^**) LMC motor neurons in the lumbar spinal cord was expressed as a percentage of total motor neurons [Foxp1**^+^** MNs (%)] (M) or electroporated motor neurons [GFP^+^ MNs (%)] (N), with 5 embryos per group. (O-V) Detection of neurofilament and GFP in the limb nerve branches of the crural plexus in chick embryos electroporated with different plasmids and siRNAs, including *GFP* (O), *scrambled [Sp1]siRNA* and *GFP* (P), *[Sp1]siRNA* and *GFP* (Q), *Sp1* and *GFP* (R), *[Sp1]siRNA, mouse Sp1* and *GFP* (S), *Sp1^T278A, T739A^* and *GFP* (T), medial LMC axonal marker, *e[Isl1]::GFP* (U), or *[Sp1]siRNA* and *e[Isl1]::GFP* (V). GFP fluorescence signals were quantified in dorsal and ventral limb nerves, with the percentages (GFP Fluo [%]) presented for each electroporation condition. The number of embryos was **n = 5** for all groups. d, dorsal; v, ventral; n.s. = not significant; *** = p<0.001; ** = p<0.01; * = p<0.05; statistical significance computed using Fisher’s exact test (A-N) or Mann-Whitney U test (O-U); solid lines in each scatterplot show the median difference. Scale bar: (A-C, H-J) 45 μm; (D-F) 35μm; (O-U) 150 μm

To assess whether Sp1 knockdown influences the trajectory selection of LMC axons within the limb, we quantified the proportions of GFP^+^ axons in the dorsal and ventral limb nerve branches by measuring the fluorescence intensities across a series of hindlimb section images from multiple embryos under each experimental condition (Kania and Jessell, 2003; Luria et al., 2008). For statistical analysis, we compared the proportions of GFP^+^ axons projecting dorsally or ventrally between the different experimental conditions. In embryos co-electroporated with *[Sp1]siRNA* and *GFP*, a significantly greater proportion of GFP^+^ axons was found in the ventral nerve branches compared to embryos electroporated with either the *GFP* or *scrambled [Sp1]siRNA* controls (Fig. 2O-Q). Notably, the axon misrouting effect was corrected by co-expressing mouse *Sp1* (Fig. 2Q, S). These results suggest that Sp1 knockdown leads to a significantly higher proportion of LMC motor axons projecting into the ventral limb.

We then conducted Sp1 gain-of-function experiments by co-electroporating *Sp1* and *GFP* expression plasmids into LMC neurons and examined motor axon trajectories in the limb. Compared to *GFP* controls, the axon length, specification, and survival of LMC neurons were unaffected in embryos co-expressing *Sp1* and *GFP* (Fig. 2H, I, K-M), and the proportion of electroporated GFP^+^ LMC neurons was consistent across both LMC divisions (Fig. 2H, I, N). In embryos co-expressing *Sp1* and *GFP*, a significantly higher proportion of GFP^+^ axons was observed in the dorsal nerves compared to *GFP* controls (Fig. 2O, R), suggesting that Sp1 overexpression increases the number of LMC motor axons entering the dorsal limb. To further explore the role of Sp1’s regulatory activity, we used a *Sp1^T278A, T739A^* expression plasmid, which encodes a mutant form of Sp1 that cannot be phosphorylated (Chuang et al., 2008). In comparison to *GFP* controls, LMC axon outgrowth, neuron specification, and survival remained normal in embryos co-expressing *Sp1^T278A, T739A^*, and *GFP* (Fig. 2J-N). However, similar to the effects of Sp1 knockdown, co-expression of *Sp1^T278A, T739A^*, and *GFP* resulted in a significantly higher proportion of GFP^+^ axons in the ventral nerve branches compared to *GFP*-only controls (Fig. 2O, T). Collectively, these results demonstrate that Sp1 expression in LMC motor neurons is crucial for accurate axon guidance in the limb.

Two potential explanations could account for the increased proportion of LMC neurons projecting into the ventral nerve branch after Sp1 knockdown: 1) some lateral LMC axons may enter the ventral limb nerve, or 2) both medial and lateral LMC axons project into both limb nerves, but a greater proportion of lateral LMC axons enter the ventral limb nerve. Regardless, Sp1 knockdown clearly results in a loss of precision in the trajectory selection of lateral LMC axons. To investigate whether Sp1 knockdown leads to medial LMC axons being misdirected into the dorsal limb mesenchyme, we co-electroporated *[Sp1]siRNA* with the *e[Isl1]::GFP* plasmid, which preferentially labels medial LMC motor neurons and their axons (Kao et al., 2009), and used *e[Isl1]::GFP* alone as a control. In embryos co-electroporated with *[Sp1]siRNA* and *e[Isl1]::GFP*, a significantly higher proportion of GFP^+^ axons were observed in the dorsal limb nerve compared to the *e[Isl1]::GFP* control group (Fig. 2T, U). Collectively, these results demonstrate that Sp1 is essential for ensuring accurate limb trajectory selection by both lateral and medial LMC axons.

### Asymmetric Sp1 knockdown sensitivity of the LMC trajectory choice

To determine the extent to which both medial and lateral LMC divisions are affected by Sp1 loss of function, we analyzed LMC axon trajectory in *Sp1* mutant mice. Since complete deletion of Sp1 results in lethality by e10.5, before LMC axons extend into the limb at e11.5-12.5 (Marin et al., 1997), we used mice with neuron-specific Sp1 deletion (*Syn1-Cre:Sp1^F/F^*) to study Sp1’s role in LMC trajectory selection (Zhu et al., 2001). In *Syn1-Cre:Sp1^F/F^*mice, LMC neuron specification and survival were comparable to those in Sp1 floxed (*Sp1^F/F^*) littermate controls (Fig. 3A-J). To trace LMC axon trajectories, we labeled LMC neurons using HRP retrograde tracer injections into the dorsal or ventral shank muscles of e12.5 *Syn1-Cre:Sp1^F/F^*and *Sp1^F/F^* littermate embryos and quantified the proportions of tracer-filled LMC neurons expressing Isl1 and Lim1 (Kania et al., 2000; Chang et al., 2018). In *Syn1-Cre:Sp1^F/F^* embryos, the proportion of medial LMC neurons labeled by dorsal limb HRP injection was significantly higher compared to control embryos (Fig. 3K-O). Similarly, a significantly greater proportion of lateral LMC neurons was labeled by ventral limb HRP injection in *Syn1-Cre:Sp1^F/F^* embryos than in control embryos (Fig. 3Q-U). These results indicate that Sp1 is essential for the proper selection of LMC axon trajectories and suggest that medial and lateral LMC neurons differ in their reliance on Sp1 function (Fig. 3P, V).

**Figure 3.**
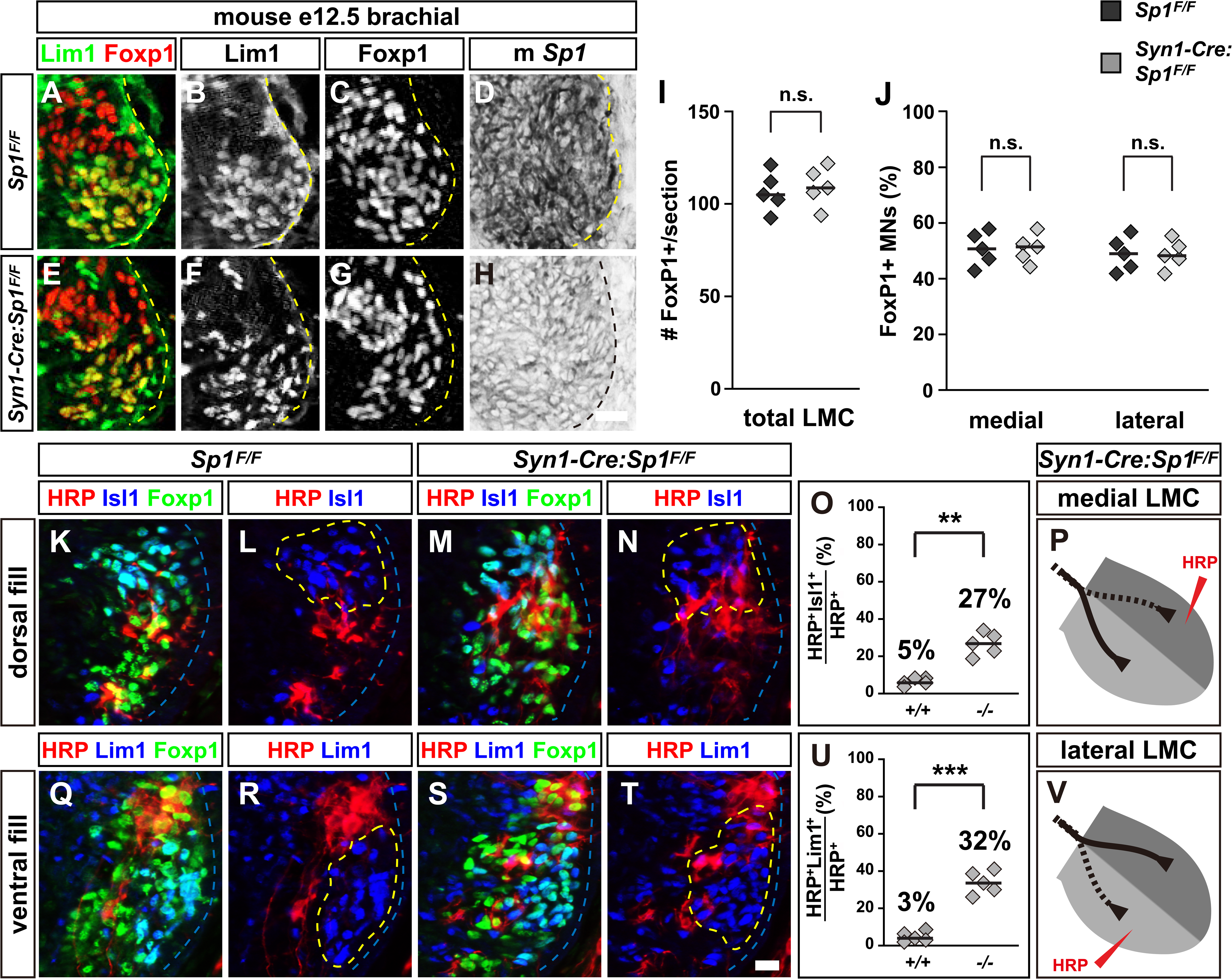
Sp1 is required for the fidelity of LMC motor axon trajectory selection. Retrograde labeling of LMC neurons via HRP injections into dorsal or ventral limb muscles of mouse embryos at e12.5. (A-H) Detection of Lim1 (green), Foxp1 (red), and Sp1 in the LMC region at the brachial level in mouse *Sp1^F/F^* (A-D) or *Syn1-Cre:Sp1^F/F^*(E-H) embryos. (I) Quantification of LMC motor neurons as the average number of total (Foxp1^+^) LMC neurons per section (# FoxP1^+^/section). The number of embryos used: n = 5 for all groups. (J) Quantification of total medial (FoxP1^+^ Lim1^-^) and lateral (FoxP1^+^ Lim1^+^) LMC motor neurons in the brachial spinal cord, expressed as a percentage of total motor neurons [FoxP1^+^ MNs (%)]. The number of embryos used: n = 5 for all groups. (K-N) Detection of HRP (red), Isl1 (blue), and Foxp1 (green) in the LMC regions of *Sp1^F/F^* (K, L) and *Syn1-Cre:Sp1^F/F^*(M, N) embryos following HRP injection into the dorsal forelimb muscles. (O) Quantification of retrogradely labeled medial LMC axon projections. The graph displays the percentage of HRP^+^ motor neurons expressing the medial LMC marker Isl1 after dorsal limb injection. The number of embryos used: n = 5 for all groups. (P) Schematic summary showing medial LMC projections in *Syn1-Cre:Sp1^F/F^* mice, highlighting the significant misrouting of medial LMC axons into the dorsal limb. (Q-T) Detection of HRP (red), Lim1 (blue), and Foxp1 (green) in the LMC regions of *Sp1^F/F^* (Q, R) and *Syn1-Cre:Sp1^F/F^*(S, T) embryos following HRP injection into the ventral forelimb muscles. (U) Quantification of retrogradely labeled lateral LMC axon projections. The graph displays the percentage of HRP^+^ motor neurons expressing the lateral LMC marker Lim1 after ventral limb injection. The number of embryos used: n = 5 for all groups. (V) Schematic summary showing lateral LMC projections in *Syn1-Cre:Sp1^F/F^* mice, depicting significant misrouting of lateral LMC axons into the ventral limb. HRP, horseradish peroxidase; *Syn1*, *Synapsin1*; n.s. = not significant; ** = p<0.01; *** = p<0.001; statistical significance computed using Fisher’s exact test; solid lines in each scatterplot show the median difference. Scale bar: (A-H, K-N, Q-T) 20 μm

### Sp1 regulates axonal guidance and neuronal development via ephrin signaling

To further determine Sp1 function in neural development, RNA sequencing was performed to elucidate the role of Sp1 in regulating gene expression during neuronal development by comparing *Syn1-Cre:Sp1^F/F^*neurons from to their control counterparts. The analysis revealed significant changes in gene expression, as shown in the volcano plot, with 333 genes upregulated and 353 genes downregulated in response to Sp1 modulation (Fig. 4A). This indicates that Sp1 plays a broad and essential role in neuronal gene regulation. To understand the functional implications of these differentially expressed genes, pathway enrichment analysis using Ingenuity Pathway Analysis was conducted (Fig. 4B). The results pointed to key developmental processes, particularly highlighting pathways involved in axonal guidance and ephrin-receptor signaling, suggesting that Sp1 influences critical signaling pathways that guide neuronal connectivity and axon navigation. The network analysis of these pathways further provided insights into the molecular interactions, where transcription factors, receptors, and enzymes were mapped in relation to Sp1 activity (Fig. 4C). Upregulated genes were primarily involved in enhancing neuron signaling, while downregulated genes suggested suppression of specific pathways. Among these, genes like *ephrin-B2 (Efnb2)* and *ephrin-A5 (Efna5)*, both of which play pivotal roles in axon guidance, were significantly downregulated following Sp1 knockdown (Fig. 4D). Conversely, *Sema7A*, another gene involved in neuronal guidance, showed increased expression, implying that Sp1 may serve as both a promoter and repressor, depending on the target gene and context (Fig. 4D).

**Figure 4.**
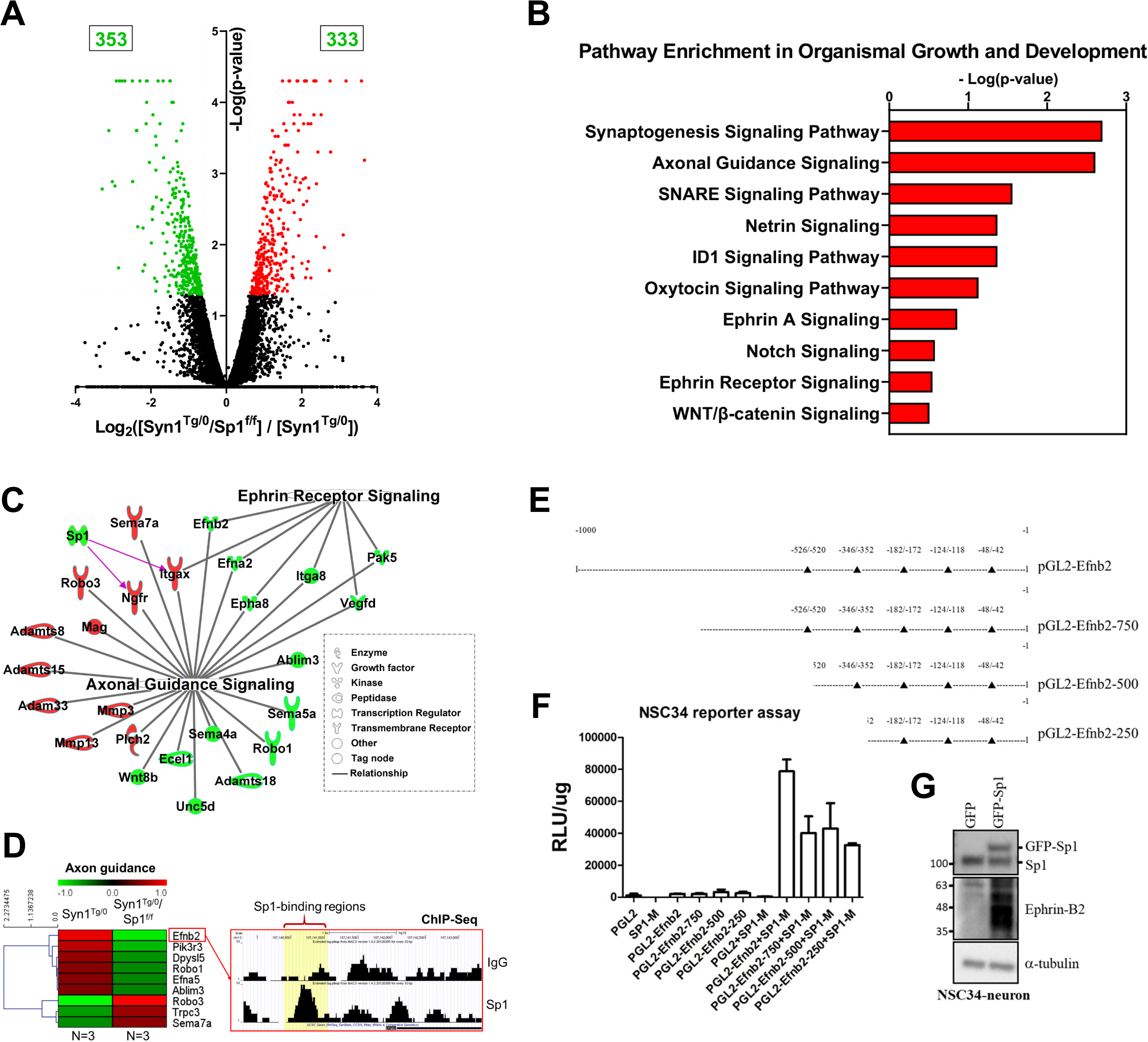
Sp1-mediated gene expression and pathways analysis in neuronal development. **(A)** Volcano plot showing the differential gene expression between *Syn1^Tg/0^/Sp1^f/f^*and *Syn1^Tg/0^* conditions. The x-axis represents the Log2 fold-change in gene expression, and the y-axis shows the −Log (p-value). Significantly upregulated genes (in red) and downregulated genes (in green) are displayed, with 333 and 353 genes identified as upregulated and downregulated, respectively. (B) Pathway enrichment analysis was performed on the differentially expressed genes (DEGs) using Ingenuity Pathway Analysis, focusing on pathways related to organismal growth and development. The top 10 enriched pathways and signaling events are presented. (C) Network representation of key pathways, including Ephrin-Receptor Signaling and Axonal Guidance Signaling. DEGs are represented, with red nodes indicating upregulated genes and green nodes showing downregulated genes. The network highlights the interactions between these genes, with different node shapes corresponding to the functional classes of the gene products (e.g., transcription factors, receptors, enzymes). (D) RNA sequencing analysis of gene expression in primary mouse neurons revealed significant changes in genes related to axon guidance. Specifically, *ephrin-B2 (Efnb2)* and *ephrin-A5 (Efna5)* expression decreased, while *Sema7A* increased following Sp1 siRNA knockdown (left panel). A ChIP assay using Sp1 antibody confirmed Sp1 binding at the ephrin-B2 promoter region (right panel). (E-G) Diagrams illustrate the predicted Sp1-binding sites on the *ephrin-B2* promoter (E). Both promoter activation and protein levels were assessed in NSC-34 cells, indicating Sp1’s regulatory role (F, G).

To investigate whether Sp1 directly regulates these axon guidance-related genes, a ChIP assay was performed, which confirmed that Sp1 binds directly to the *ephrin-B2* promoter, suggesting transcriptional control (Fig. 4E). This was further supported by NSC-34 cell experiments, where knockdown of Sp1 led to a notable reduction in both *ephrin-B2* promoter activity and protein levels, thereby confirming Sp1’s role as a positive regulator of ephrin-B2 (Fig. 4F, G). These findings imply that Sp1 exerts a precise regulatory influence on genes involved in axonal navigation, either by directly enhancing or repressing their transcription, ultimately affecting neuronal network formation. Together, these data suggest the potential role of Sp1 to mediate a range of gene expression programs crucial for neuronal development, especially in the context of axon guidance and synaptic signaling.

### Sp1 is specifically required for ephrin-mediated motor axon responses

To explore the role of Sp1 in Eph signaling, we assessed the response of LMC axons to ephrin and Eph protein stripes in the context of Sp1 loss of function (Kao and Kania, 2011). LMC explants were taken from HH stage 25/26 embryos and placed on alternating stripes: one containing a mix of ephrin-Fc and a Cy3 secondary antibody, and the other containing only Fc protein. Medial LMC axons were marked by *e[Isl1]::GFP* electroporation, while lateral LMC axons were identified by their EphA4 expression. Stripe preference was quantified by measuring the proportion of GFP or EphA4 signal over the respective stripes after overnight explant culture (Kao and Kania, 2011; Kao et al., 2015). Neurite growth preference in ephrin versus Fc stripes was compared statistically between different experimental conditions.

Medial LMC neurons co-electroporated with *[Sp1]siRNA* and *e[Isl1]::GFP* showed a significantly increased repulsion from ephrin-A5 stripes compared to controls expressing only *e[Isl1]::GFP*. This indicates that Sp1 influences ephrin-A expression, ensuring ephrin-A/EphA *cis*-attenuation in medial LMC axons (Fig. 5A, B). Additionally, medial LMC neurons co-electroporated with *[Sp1]siRNA* and *e[Isl1]::GFP* exhibited significantly enhanced attraction toward EphA4 stripes compared to controls, suggesting possible EphA:ephrin-A reverse signaling, potentially due to reduced ephrin-A levels following Sp1 knockdown (Fig. 5C, D). Given that previous studies have identified Netrin-1 and Semaphorin (Sema) 3F as additional regulators of medial LMC axon guidance into the limb (Poliak et al., 2015; Huber et al., 2005; Moret et al., 2007), we also tested medial LMC axon responses to Netrin-1 and Sema3F stripes under Sp1 knockdown conditions. Medial LMC neurons with Sp1 knockdown showed normal avoidance of Netrin-1 and Sema3F stripes, similar to controls expressing *e[Isl1]::GFP* alone (Fig. 5E-H), indicating that Sp1 does not play a significant role in Netrin-1- or Sema3F-mediated medial LMC axon guidance *in vitro*.

**Figure 5.**
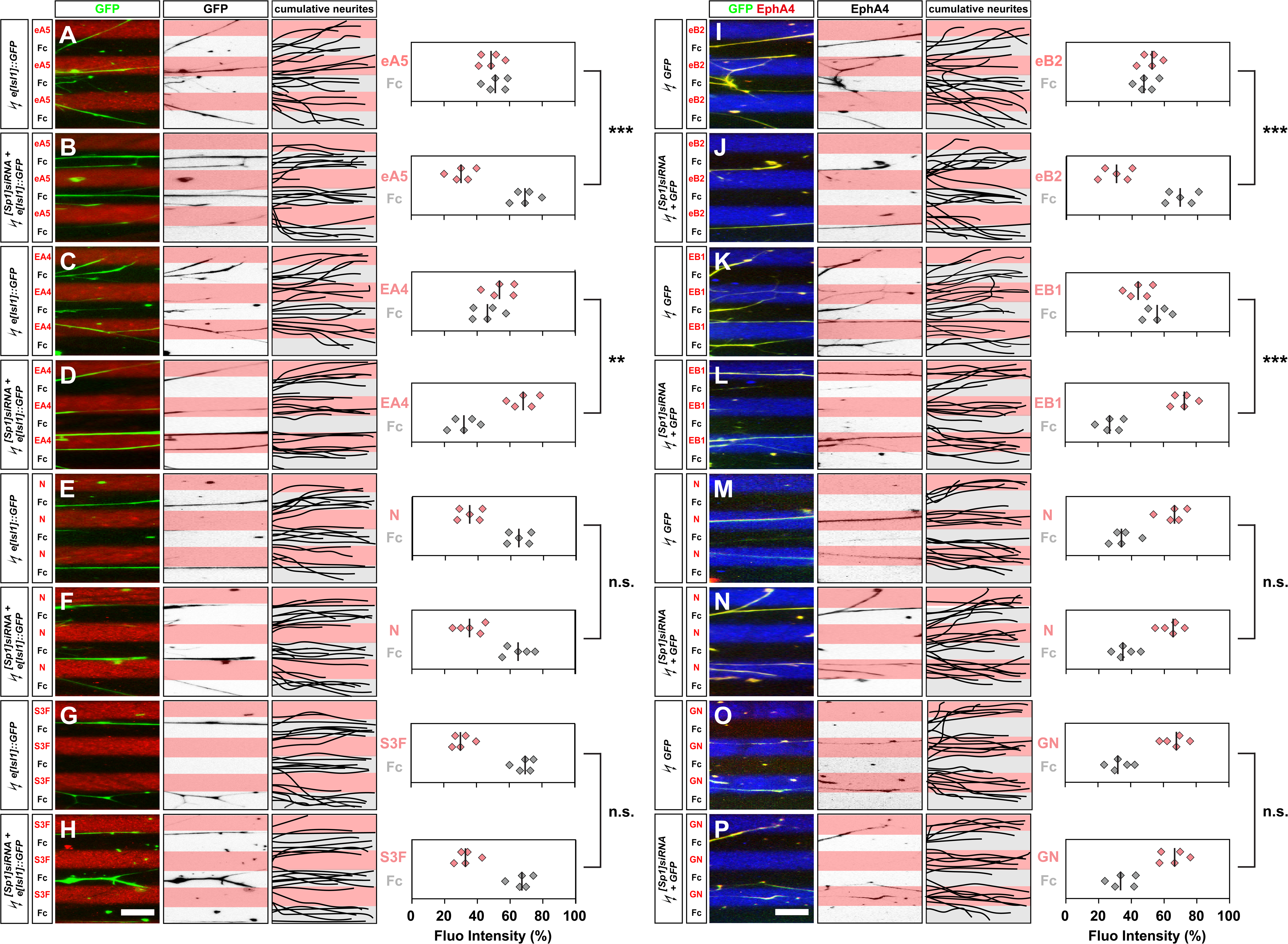
Sp1 function is specifically required for ephrin-mediated guidance of cultured LMC neurites. Growth preference of medial or lateral LMC axons on protein stripes. Each experiment consists of three panels (left, middle, and right) and one quantification. (A-H) Left panels: Detection of medial (GFP^+^) LMC neurites from explants grown on stripes of eA5/Fc (A), EA4/Fc (C), N/Fc (E), or S3F/Fc (G), and explants co-electroporated with *[Sp1]siRNA* on stripes of eA5/Fc (B), EA5/Fc (D), N/Fc (F), or S3F/Fc (H) stripes. Middle panels: Inverted images showing GFP signals as dark pixels overlaid on the substrate stripes. Right panels: Superimposed images of five representative explants from each experimental group, illustrating the distribution of medial LMC neurites. Quantification shows medial (GFP^+^) LMC neurites on the first (pink) and second (pale) stripes as a percentage of total GFP signals. A minimum of 88 neurites was analyzed. Number of embryos: n = 5 for all groups. A minimum of 3 explants per embryo was used. (I-P) Left panels: Detection of lateral (GFP^+^ EphA4^+^) LMC neurites from explants grown on stripes of eB2/Fc (I), EB1/Fc (K), N/Fc (M), or GN/Fc (O), and explants co-electroporated with *[Sp1]siRNA* on stripes of eB2/Fc (J), EB1/Fc (L), N/Fc (N), or GN/Fc (P). Middle panels: Inverted images showing EphA4 signals as dark pixels overlaid on the substrate stripes. Right panels: Superimposed images of five representative explants from each experimental group, demonstrating the distribution of lateral LMC neurites. Quantification shows lateral (EphA4^+^) LMC neurites on the first (pink) and second (pale) stripes as a percentage of total EphA4 signals. A minimum of 85 neurites was analyzed. Number of embryos: **n = 5** for all groups. A minimum of 3 explants per embryo was used. eA5, ephrin-A5-Fc; EA4, EphA4-Fc; N, Netrin-1; S3F. Semaphorin-3F; eB2, ephrin-B2-Fc; EB1, EphB1-Fc; GN, GDNF; *** = p<0.001; ** = p<0.01; n.s. = not significant; statistical significance computed using Mann-Whitney U test; solid lines in each scatterplot show the median difference. Scale bar: (A-P) 150 μm

In contrast, lateral LMC neurons co-electroporated with *[Sp1]siRNA* and *GFP* showed significantly increased repulsion from ephrin-B2 stripes compared to controls expressing only *GFP*. This indicates that Sp1 regulates ephrin-B expression to ensure ephrin-B/EphB *cis*-attenuation in lateral LMC axons (Fig. 5I, J). Furthermore, lateral LMC neurons co-electroporated with *[Sp1]siRNA* and *GFP* displayed significantly increased attraction to EphB1 stripes compared to controls, suggesting the presence of EphB:ephrin-B reverse signaling, potentially caused by reduced ephrin-B levels following Sp1 knockdown (Fig. 5K, L). Given previous findings indicating that Netrin-1 and GDNF signaling are other pathways involved in regulating lateral LMC axon trajectory into the limb, we next tested the response of lateral LMC axons to Netrin-1 and GDNF stripes. Lateral LMC neurons co-electroporated with *[Sp1]siRNA* and *GFP* exhibited normal neurite attraction toward Netrin-1 (Fig. 5M, N) and GDNF stripes (Fig. 5O, P) compared to controls expressing only *GFP*. This suggests that, similar to medial LMC axons, Sp1 does not play a critical role in Netrin-1- or GDNF-mediated lateral LMC axon guidance *in vitro*. Taken together, these findings indicate that Sp1 is specifically required for ephrin-A- and ephrin-B-mediated responses in LMC axons.

### Sp1 coactivators, p300 and CBP, are required for Sp1-regulated LMC pathfinding

Two Sp1 coactivators, p300 and CBP, are present in LMC neurons during the proper trajectory of their axons into the limb (Fig. 1). To explore the role of p300 and CBP in Sp1-regulated LMC axon pathfinding, we investigated whether their loss would reduce the enhanced growth preference of LMC neurites against ephrin stripes that occurs after Sp1 overexpression. In comparison to GFP controls, axon length, neuron specification, and survival were normal in embryos co-expressing *[p300]siRNA* (or *[CBP]siRNA*) and *GFP* (Fig. 6A-K). Similarly, GFP^+^ LMC neurons displayed comparable proportions of electroporated cells in both LMC divisions (Fig. 6L). Medial LMC neurons co-electroporated with *Sp1* and *e[Isl1]::GFP* showed significantly increased repulsion from ephrin-B2 stripes compared to controls expressing *e[Isl1]::GFP* alone (Fig. 6M, N). However, when these neurons were co-electroporated with *[p300]siRNA* and *[CBP]siRNA*, the intensified repulsion effect was eliminated compared to controls expressing *Sp1* and *e[Isl1]::GFP* only or those co-expressing *scrambled [p300]siRNA* and *scrambled [CBP]siRNA* (Fig. 6O, P), indicating that p300 and CBP are essential for Sp1’s role in ephrin-mediated medial LMC axon guidance *in vitro*.

**Figure 6.**
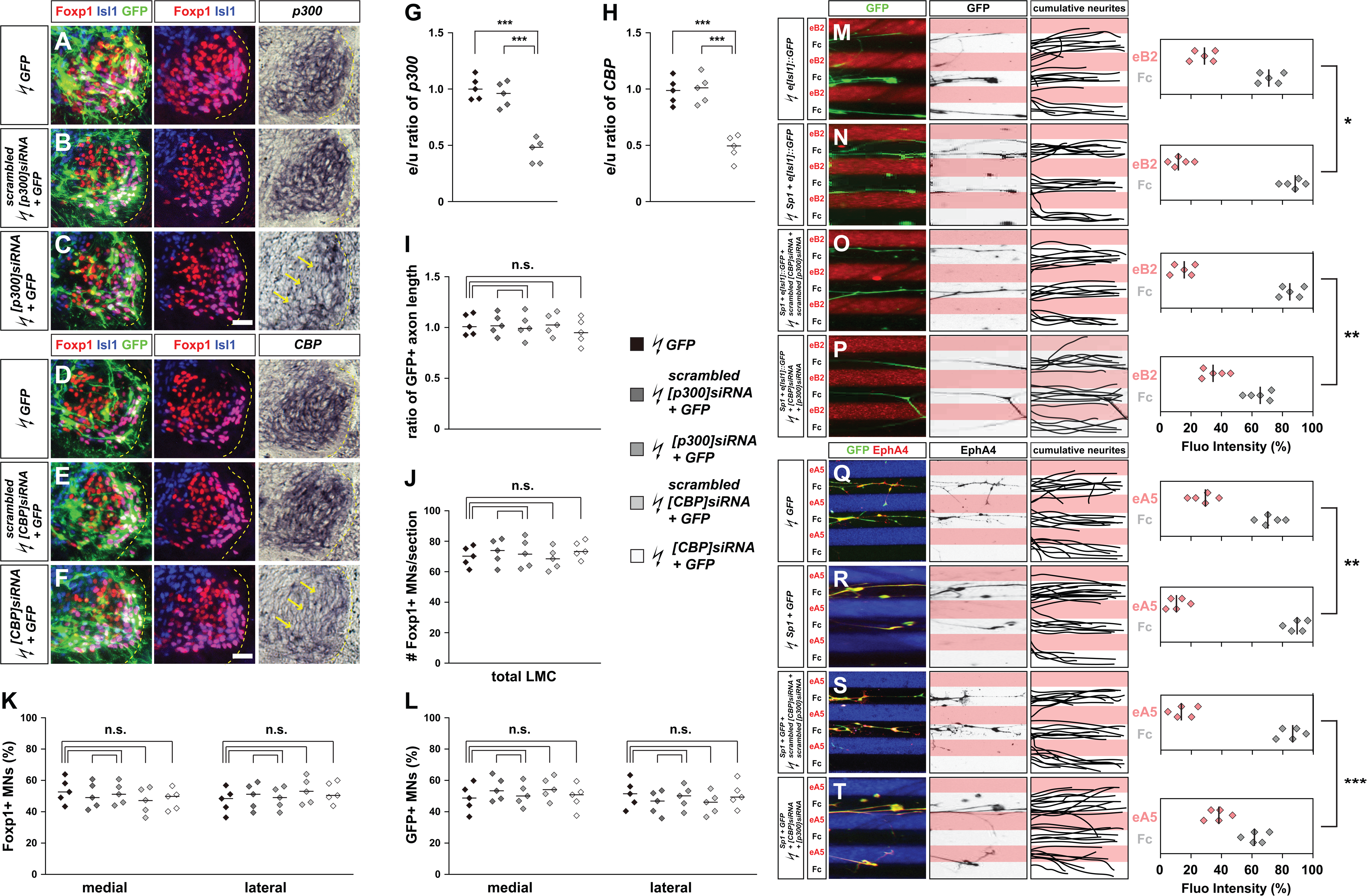
Sp1 coactivators, p300 and CBP, are required for Sp1-regulated guidance of cultured LMC neurites. (A-L) All images are from chick embryos at HH stage 28/29, lumbar levels. (A-C) Detection of Foxp1, Isl1, GFP, and p300 in LMC neurons from chick embryos electroporated with *GFP* (A), *scrambled [p300]siRNA* and *GFP* (B), or *[p300]siRNA* and *GFP* (C). (D-F) Detection of Foxp1, Isl1, GFP, and *CBP* in LMC neurons from chick embryos electroporated with *GFP* (A), *scrambled [CBP]siRNA* and *GFP* (B), or *[CBP]siRNA* and *GFP* (C). (G, H) Quantification of the effects of *GFP*, *scrambled [p300]siRNA* and *GFP*, or *[p300]siRNA* + *GFP* electroporation on *p300* mRNA levels (G) and the effects of *GFP*, *scrambled [CBP]siRNA* and *GFP*, or *[CBP]siRNA* + *GFP* electroporation on *CBP mRNA* levels (H). The ratio of immunoreactivity between electroporated and un-electroporated contralateral sides (e/u ratio) was measured from at least 15 sections of 5 embryos. (I) Quantification of the effects of *GFP*, *scrambled [p300]siRNA* and *GFP*, *[p300]siRNA* and *GFP*, *scrambled [CBP]siRNA* and *GFP*, or *[CBP]siRNA* and *GFP* electroporation on axon outgrowth. GFP^+^ LMC axon lengths were compared to the GFP control group, with measurements taken from at least 15 sections of 5 embryos. (J) The number of LMC motor neurons is expressed as the average number of total (Foxp1^+^) LMC neurons per section (**#** Foxp1+ MNs/section). Numbers of embryos: n = 5 for all groups. (K, L) The number of total or electroporated medial (Foxp1**^+^** Isl1**^+^**) and lateral (Foxp1**^+^** Isl1**^-^**) LMC motor neurons in the lumbar spinal cord is expressed as a percentage of total motor neurons [Foxp1^+^ MNs (%)] (K) or electroporated motor neurons [GFP^+^ MNs (%)] (L). Numbers of embryos: n = 5 for all groups. (M-T) Growth preferences of medial or lateral LMC axons on protein stripes. Each experiment includes three panels (left, middle, and right) and one quantification. (M-P) Left panels: Detection of medial (GFP**^+^**) LMC neurites from explants electroporated with *GFP* plasmid electroporated explants (M), *Sp1* expression plasmid co-electroporated explants (N), *Sp1* + *scrambled [CBP]siRNA* + *scrambled [p300]siRNA* (O), or *Sp1* + *[CBP]siRNA* + *[p300]siRNA* (P) on eB2/Fc stripes. Middle panels: Inverted images showing GFP signals as dark pixels overlaid on substrate stripes. Right panels: Superimposed images of five representative explants from each group, highlighting the distribution of medial LMC neurites. Quantification of medial (GFP**^+^**) LMC neurites on the first (pink) and second (pale) stripes, expressed as a percentage of total GFP signals. A minimum of 88 neurites were analyzed. Numbers of embryos: n = 5 for all groups, with a minimum of 3 explants per embryo. (Q-T) Left panels: Detection of lateral (GFP^+^ EphA4^+^) LMC neurites from explants electroporated with *GFP* plasmid (Q), *Sp1* expression plasmid (R), *Sp1* + *scrambled [CBP]siRNA* + *scrambled [p300]siRNA* (S), or *Sp1* + *[CBP]siRNA* + *[p300]siRNA* stripes. Middle panels: Inverted images showing EphA4 signals as dark pixels overlaid on substrate stripes. Right panels: Superimposed images of five representative explants from each group, highlighting the distribution of lateral LMC neurites. Quantification of lateral (EphA4**^+^**) LMC neurites on the first (pink) and second (pale) stripes, expressed as a percentage of total EphA4 signals. A minimum of 85 neurites were analyzed. Numbers of embryos: n = 5 for all groups, with a minimum of 3 explants per embryo. eB2, ephrin-B2-Fc; eA5, ephrin-A5-Fc; *** = p<0.001; ** = p<0.01; * = p<0.05; statistical significance computed using Fisher’s exact test (A-L) or Mann-Whitney U test (M-T); solid lines in each scatterplot show the median difference. Scale bar: (A-F) 45 μm; (M-T) 150 μm

Similarly, lateral LMC neurons co-electroporated with *Sp1* and *GFP* showed significantly increased repulsion from ephrin-A5 stripes compared to controls expressing *GFP* alone (Fig. 6Q, R). This repulsion was abolished when *[p300]siRNA* and *[CBP]siRNA* were co-electroporated, compared to controls co-expressing *Sp1* and *GFP* alone or those co-expressing *scrambled [p300]siRNA* and *scrambled [CBP]siRNA* (Fig. 6S, T). These findings demonstrate that Sp1 cooperates with its coactivators, p300 and CBP, to regulate ephrin gene expression and mediate LMC axon pathfinding.

### Locomotor function impairment in Sp1 mutants

To investigate the role of Sp1 in regulating motor function and neuronal connectivity, we performed a series of behavioral assays on adult *Sp1^F/F^*and *Syn1-Cre:Sp1^F/F^* mice, aiming to assess locomotor activity, cognitive function, motor coordination, and muscle strength. We first evaluated general locomotor activity and anxiety-like behavior using the open field test (Figures 7A, B). Mice were placed in an open field arena, and both total distance traveled (an indicator of overall locomotor activity) and time spent in the center (a measure of anxiety-like behavior) were recorded over 60 minutes. There were no significant differences in either total distance or center duration between *Sp1^F/F^* and *Syn1-Cre:Sp1^F/F^*groups, suggesting that the deletion of Sp1 in neurons does not affect basic locomotor behavior or induce heightened anxiety. To further assess the cognitive function of the mice, we performed the Y-maze test (Figure 7C), which evaluates spatial working memory by measuring the proportion of spontaneous alternation between the maze arms. This test is commonly used to assess short-term memory and cognitive flexibility. Our data showed no significant differences in alternation percentages between the groups, indicating that spatial memory and cognitive flexibility remain unaffected by Sp1 loss. These results suggest that the cognitive processes underlying basic exploratory behavior and working memory are preserved in Sp1 mutant mice.

**Figure 7.**
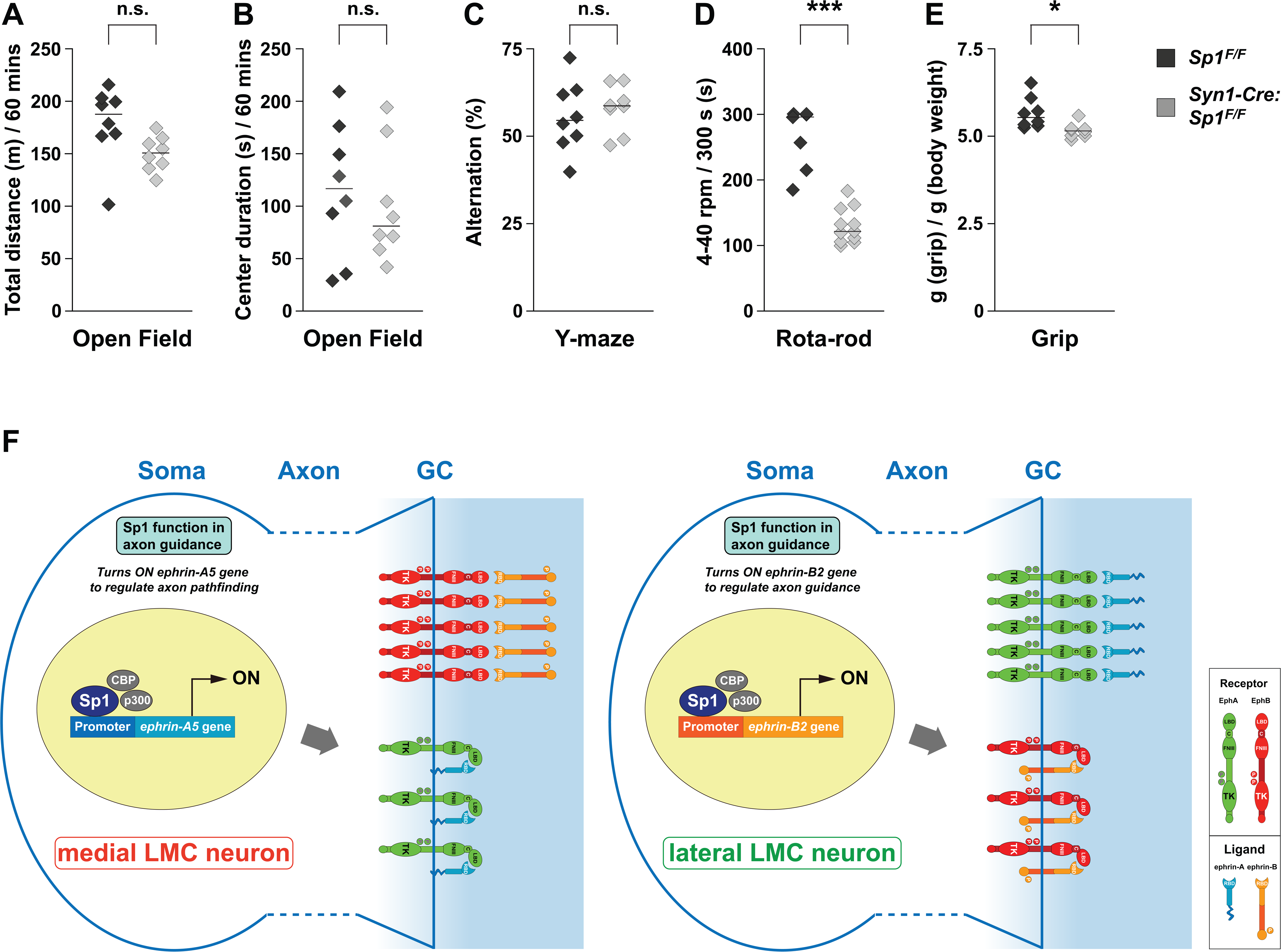
Sp1 mutant mice exhibit muscle strength and locomotor defects due to its regulation of ephrin activity in spinal motor neurons. (A, B) Open field test: The total distance traveled (A) and the time spent in the center (B) were recorded over a 60-minute period. (C) Y-maze test: The percentage of alternation between the arms was measured to assess cognitive performance. (D) Rota-rod test: Mice were tested on an accelerating rota-rod (4-40 rpm) for a maximum duration of 300 seconds. The time until the mice fell from the apparatus was recorded as a measure of motor coordination. (E) Grip strength test: Grip strength was measured using a grip strength meter, and the value was normalized to the body weight to represent the relative grip strength of the mice. (F) Sp1, in combination with its co-activators p300 and CBP, regulates the accurate trajectory of LMC axons into the limb by controlling ephrin gene expression. This regulation mediates ephrin/Eph *cis*-attenuation and ephrin:Eph *trans*-signaling, ensuring proper axon guidance. Numbers of embryos: n = 8 for both groups in open field, Y-maze, and grip tests; n = 7 for *Sp1^F/F^*, and 11 for *Syn1-Cre:Sp1^F/F^*groups in Rota-rod test.

Despite the normal performance in general locomotor and cognitive tests, *Syn1-Cre:Sp1^F/F^* mice displayed significant deficits in motor coordination and muscle strength. In the Rota-rod test (Figure 7D), which assesses motor coordination and balance, *Syn1-Cre:Sp1^F/F^* mice showed a markedly reduced latency to fall compared to controls, indicating impaired motor coordination. This suggests that Sp1 is critical for maintaining motor function, particularly in tasks that require balance and coordination. Additionally, in the grip strength test (Figure 7E), *Syn1-Cre:Sp1^F/F^* mice exhibited significantly reduced grip strength compared to controls when normalized to body weight. This decrease in muscle strength highlights that the loss of Sp1 affects not only coordination but also the physical strength of motor output.

Taken together, the combination of our *in vivo* and *in vitro* studies demonstrates that while Sp1 deficiency disrupts motor coordination and muscle strength, which are likely due to aberrant ephrin/Eph signaling in spinal motor neurons, caused by the loss of Sp1-mediated transcriptional regulation of ephrin genes. Therefore, Sp1 plays a critical role in maintaining proper motor function through its regulation of axon guidance pathways in the spinal cord (Fig. 7F).

## Discussion

Sp1 has been proposed to regulate several genes essential for early neuronal differentiation, axon growth, and synaptogenesis. Consequently, we explored its potential role in axon pathfinding *in vivo* and found that Sp1 is crucial for the accurate selection of motor axon trajectories in the limb. In this context, we discuss Sp1’s involvement in axon outgrowth, neuronal survival, differentiation, the role of CBP/p300 as coactivators in Sp1-mediated pathfinding within the LMC, and its implications for ephrin/Eph signaling.

### Sp1 is critical for motor axon guidance but is not required for axon outgrowth, neuronal survival, or migration

Sp1 participates in numerous cellular functions, including neuronal development, axon outgrowth, survival, and differentiation. Research has shown that Sp1 plays diverse roles in both early and late stages of neuronal development, particularly in axon differentiation and growth, which are crucial for establishing neuronal connections (Okamoto et al., 2002; Billon et al., 1999; Mondanizadeh et al., 2015; Hung et al., 2020). For axon outgrowth, Sp1 is thought to regulate downstream target genes essential for cytoskeletal organization and neuronal signaling. Studies indicate that Sp1 controls the expression of GAP-43 (Growth-Associated Protein 43), crucial for axonal growth and regeneration, and Tau, which stabilizes microtubule structures in developing axons (Kawasaki et al., 2023; Heicklen-Klein et al., 2000; Santpere et al., 2006; Villa et al., 2013). Additionally, Sp1 regulates the expression of axon guidance molecules like Semaphorin receptors and ephrins, which direct axon navigation through interactions with external signals (Rossignol et al., 2003; Sohl et al., 2010; Arvanitis and Davy, 2012). Sp1 is also implicated in the regulation of ROCK1, a kinase that modulates cytoskeletal reorganization.

Studies suggest that Sp1 enhances ROCK1 expression, promoting axonal extension by controlling actin filament formation in neurons (Zhang et al., 2016; Wang et al., 2020). Interestingly, we did not observe changes in limb nerve axon outgrowth in chick embryos with altered Sp1 expression in LMC neurons. Although residual Sp1 expression after siRNA knockdown may be sufficient for axon outgrowth, Sp1 appears to have a limited impact on motor axon outgrowth. This may be due to different neuronal types expressing distinct repertoires of Sp1 targets and interacting with unique kinetics and affinities.

Beyond its role in axon outgrowth, Sp1 is also thought to play a part in neuronal differentiation and survival. Research has shown that Sp1 regulates neurotrophic factors such as brain-derived neurotrophic factor (BDNF), whose expression is essential for proper neuronal differentiation into mature forms (Guida et al., 2017). Their study found that the loss of Sp1 in neuronal cultures led to reduced BDNF expression and hindered neuronal differentiation, suggesting a role for Sp1 in determining neuronal fate. Additionally, Sp1 is implicated in regulating genes involved in synaptogenesis, such as synapsin I, which is vital for synaptic vesicle transport and neurotransmitter release. Sp1 promotes synapsin I expression, indicating its function in synapse maturation and activity during differentiation (Paonessa et al., 2013).

Collectively, these studies imply that Sp1 influences not only early neuronal differentiation but also synaptic development and connectivity. Furthermore, Sp1 is crucial for neuronal survival, especially under stress conditions (Ryu et al., 2003; Yeh et al., 2011). Studies have shown that Sp1 and Sp3, which are activated by oxidative stress, promote neuronal survival (Ryu et al., 2003). Sp1 is also reported to regulate Bcl-2 transcription from the proximal P2 promoter in neurons, supporting its role in promoting survival (Smith et al., 1998). Additionally, Sp1 influences the expression of p53, a well-known tumor suppressor involved in apoptosis. By repressing p53 activity in neurons, Sp1 provides a mechanism to support neuronal survival in conditions that might otherwise lead to cell death (Vousden et al., 2009; Beckerman et al., 2010; Levine et al., 2009). This delicate balance between enhancing survival factors and suppressing apoptotic genes is crucial for neuronal longevity. However, we did not observe noticeable changes in the number or positioning of LMC neurons when Sp1 expression was altered in chick or mouse embryos. This suggests that Sp1’s role in cell survival may be redundant, at least in motor neurons, and that its role in cell differentiation might be limited or stage-specific. Further investigation into the expression patterns of other Sp1 targets during different stages of spinal motor neuron differentiation may help clarify these points.

### The necessity of CBP/p300 coactivators in Sp1-mediated LMC axon guidance is evident

Sp1 is expressed in most LMC neurons and is essential for correct axon trajectory in both medial and lateral LMC divisions, indicating a unique role in axon pathfinding within the LMC. This observation raises the question of how Sp1 influences guidance signaling through gene regulation. Research suggests that Sp1 regulates gene expression in neurons by interacting with coactivators such as CBP (CREB-binding protein) and p300 (Formisano et al., 2015; Toch et al., 2019; Lipinski et al., 2019). Sp1 is critical for activating the transcription of numerous neuronal genes, particularly those associated with synaptic plasticity and neuronal differentiation (Li et al., 2010; Safe et al., 2005). Additionally, Sp1 and its coactivators play roles in neuroprotection, mainly through the regulation of antioxidant genes. For example, studies have demonstrated that Sp1 upregulates antioxidant proteins like superoxide dismutase (SOD) in response to oxidative stress (Seo et al., 1996; Chang et al., 2017). Mutations or disruptions in Sp1 or its coactivator interactions can compromise these protective mechanisms, leading to increased neuronal vulnerability and degeneration, as seen in models of Huntington’s disease (Qiu et al., 2006; Steffan et al., 2000). Moreover, CBP and p300 are essential for maintaining synaptic plasticity, a key factor in learning and memory (Muller et al., 2022). Behavioral studies on CBP mutant mice also show deficits such as reduced locomotor habituation and impaired object recognition memory (Valor et al., 2011).

Overall, the interactions of Sp1 with CBP and p300 are fundamental to numerous neuronal functions, from development and plasticity to stress protection, highlighting the importance of this regulatory network in maintaining brain health. Disruptions in Sp1 or its coactivators not only impair normal neuronal function but also play a role in neurodegenerative disease pathogenesis, making Sp1 and its coactivators promising therapeutic targets. However, the exact role of CBP and p300 in Sp1-mediated axon guidance during neural circuit formation remains unclear. A possible model is that Sp1, together with CBP and p300, regulates the expression of guidance receptors or ligands, altering the ratio of neuron-expressed receptors to ligands, which in turn influences their interaction mode. Our findings, showing the loss of ephrin-mediated attenuation of ephrin/Eph signaling when Sp1 is depleted, along with the effects of non-phosphorylatable Sp1 and CBP/p300 loss on Sp1-mediated redirection, indicate that ephrin/Eph signaling may be a target of the Sp1/CBP/p300 regulatory complex in this context.

### Sp1 function in ephrin/Eph signaling

Our findings indicate that although Sp1 is expressed in most LMC neurons, its role in ensuring accurate LMC axon trajectory selection appears to vary, with lateral LMC axon guidance depending more heavily on Sp1 than medial LMC pathfinding. This differential requirement for Sp1 may arise from the need for distinct pathways in guiding each LMC division. Strong evidence supports the involvement of ephrin-A:EphA and ephrin-B:EphB forward signaling in guiding lateral and medial LMC axons, respectively (Kania and Jessell, 2003; Luria et al., 2008; Marquardt et al., 2005; Dudanova et al., 2012). Additionally, studies suggest that Eph:ephrin reverse signaling also plays a role in LMC axon pathfinding (Marquardt et al., 2005; Dudanova et al., 2012). The varying sensitivity of lateral and medial LMC neurons to Sp1 might thus reflect a differing requirement for Sp1 in the transcription of ephrin-A and ephrin-B genes. Previous research also identifies Netrin-1 as a bifunctional ligand, attracting lateral LMC axons while repelling medial ones (Poliak et al., 2015). Furthermore, other systems, such as GDNF and Semaphorin3F, are proposed to influence the correct trajectory of motor axons from LMC subpopulations (Huber et al., 2005; Moret et al., 2007; Kramer et al., 2006; Dudanova et al., 2010). Although there is limited biochemical evidence linking Sp1 with receptors like Dcc, Unc5c, c-Ret, or Neuropilin2, our *in vitro* data—showing no significant change in LMC neurite growth preferences towards Netrin-1, Sem3F, or GDNF stripes following Sp1 knockdown—suggests that Sp1’s role in Netrin-1, Sema3F, or GDNF-mediated LMC pathfinding is minimal, highlighting the specificity of Sp1’s function in regulating Eph/ephrin signaling in this context.

## Conclusion

In this study, we demonstrated that Sp1 is essential for motor axon trajectory selection *in vivo*. This is the first report identifying a significant role for Sp1 in Eph/ephrin signaling and directly comparing its involvement in ephrin-A-versus ephrin-B-mediated motor axon guidance. Our findings reveal the necessity of Sp1 in distinct Eph/ephrin pathways, underscoring the value of the LMC motor axon projection system as a model for studying simple axon guidance decisions and the function of the ephrin signaling machinery.

## Materials and Methods

### Animals

Fertilized chick eggs (JD-SPF Biotech, Miaoli, Taiwan) were stored at 18°C for no longer than one week and incubated at 38°C, with developmental stages determined according to standard protocols (Hamburger and Hamilton, 1951). *B6.Cg-Tg(Syn1-cre)671Jxm/J* mice (*Syn1-cre*) were obtained from The Jackson Laboratory (Bar Harbor, ME, USA), while *C57BL/6- Sp1tm1(GEMMS)Narl* mice (*Sp1^F/F^*) were provided by the National Laboratory Animal Center (NLAC) in Tainan, Taiwan. All mice were bred and housed at the NLAC until they reached 8 weeks of age. The genotypes of *Syn1-cre:Sp1^F/F^* mice were confirmed by PCR, using the following primers: Internal control (Forward): CTAGGCCACAGAATTGAAAGATCT; Internal control (Reverse): GTAGGTGGAAATTCTAGCATCATCC; Syn1-cre (Forward): CTCAGCGCTGCCTCAGTCT; Syn1-cre (Reverse): GCATCGACCGGTAATGCA; Sp1 (Forward): AACAGGGGCTGGAACAATAG; Sp1flox/flox (Reverse): CGTGAGTTCAAGGGAAGACTGG. All mice were housed in an air-conditioned vivarium with ad libitum access to food and water, under a 12-hour light/dark cycle. All animal handling protocols, including for chick and mouse embryos, were approved by the Institutional Animal Care and Use Committee at Taipei Medical University (Approval Number: LAC-2020-0549).

### Molecular biology

The characterization of expression constructs, including *e[Isl1]::GFP*, *Sp1*, and *Sp1^T278A, T739A^*, has been previously described (Kao et al., 2009; Chuang et al., 2008).

*In situ* hybridization cDNA probes were prepared as outlined in Kao et al. (2009). Briefly, target sequence amplification primers were designed using Primer3 version 0.4.0 software (Rozen and Skaletsky, 2000), with the probe size set between 600-800 bp. One-step RT-PCR was performed using the designed primers containing T7 polymerase promoters (Invitrogen) to amplify the cDNA template from chick HH stage 25/26 or mouse embryonic day 11.5 (e11.5) pooled brain RNA. The PCR products were purified through gel electrophoresis using a 1% agarose gel, followed by extraction using the QIAquick Gel Extraction Kit (Qiagen). The purified DNA was re-amplified via PCR, and the DNA yield was estimated using a Low DNA Mass Ladder (Invitrogen) after gel electrophoresis. DIG-labeled RNA probes were synthesized by *in vitro* transcription with T7 RNA polymerase, using the DIG RNA Labeling Kit (Roche). All probes were verified by sequencing. The source and recognized region for each probe are as follows: chick *Sp1*: NM_204604, 3943-4618; chick *p300*: XM_004937710, 4781-5484; chick *CBP*: XM_015294627, 3151-3836; mouse *Sp1*: NM_013672, 1270-1942; mouse *p300*: NM_177821, 4063-4773; mouse *CBP*: NM_001025432, 1279-2047. Details regarding *Isl1* and *Lim1* probes were previously described in Tsuchida et al. (1994).

### Chick in ovo electroporation

Spinal cord electroporation of expression plasmids was performed at HH stage 18/19 in embryos of either sex, as previously described (Momose et al., 1999; Kao et al., 2009; Luria et al., 2008). Briefly, a 5-8 µg/µL solution of plasmid DNA in TE buffer (pH 7.5, 10 mM Tris-HCl and 1 mM EDTA, Invitrogen) was injected into the lumbar neural tube through a small window created in the eggshell under a stereomicroscope (SAGE Vision). The lower bodies of the chick embryos were electroporated using platinum/iridium electrodes (FHC) and the TSS20 electroporator (Ovodyne) with the following settings: 30 V, 5 pulses, 50 ms pulse width, at 1-second intervals. The eggshell windows were sealed with Parafilm (Pechinery Plastic Packaging) and the embryos were incubated at 38°C until they were harvested at HH stage 28/29. The efficiency of electroporation varied between 5-30% of total LMC neurons, depending on the plasmid construct size and DNA concentration used.

When untagged expression plasmids were electroporated alongside *GFP* expression plasmids or *GFP*-fused plasmids, their concentration was at least three times higher than that of *GFP*-fused plasmids to ensure efficient co-expression. SiRNA duplex oligonucleotides with 3’ TT overhangs were purified using MicroSpin G-25 columns (GE Healthcare) in TE buffer containing 20 mM NaCl (Sigma). A 1 µg/µL GFP expression plasmid was co-electroporated with the siRNA solution to label motor axons. The sequences of the siRNA sense strands are as follows: *[Sp1]siRNA*, 1:1 mixture of GCAACATCACCTTGCTGCCTGTCAA and CCAAGCACATCAAGACCCACCAGAA; *scrambled [Sp1]siRNA*, 1:1 mixture of GCAACTATTCCGTCGGTCCTCACAA and CCACACCTAGAACCACACACAGGAA; *[p300]siRNA*, 1:1 mixture of CCTGGCACAAGATGCTGCCTCTAAA and TGGCCGAGGTGTTTGAGCAAGAAAT; *scrambled [p300]siRNA*, 1:1 mixture of CCTCACGAAGTAGTCTCCTCGGAAA and TGGGAGGTGTTTGAGCAAGACCAAT; *[CBP]siRNA*, 1:1 mixture of TCTCCGCCAGCGATAGCACAGATTT and CAACGTGCTGGAAGAAAGCATTAAA; *scrambled [CBP]siRNA*, 1:1 mixture of TCTACCGAGCGCGATGACAACCTTT and CAATCGTAAGGAAAGTACGTGCAAA.

### HRP retrograde labeling of LMC motor neurons

Retrograde labeling of mouse motor neurons was carried out as previously described (Luria et al., 2008; Poliak et al., 2015). Briefly, e12.5 embryos of either sex were dissected at the thoracic level and incubated at room temperature in aerated DMEM/F12 medium (Invitrogen). A 20% solution of HRP (Roche) was used as the retrograde tracer, prepared by dissolving 100 mg of HRP in 450 µL of PBS and adding 50 µL of 10% lysophosphatidylcholine (Fluka) in PBS. The HRP solution was injected into either the dorsal or ventral forelimb proximal muscle group. Following injection, the embryos were incubated at 32°C under an infrared lamp, with continuous aeration using 95% air and 5% CO_2_. Fresh medium was added every 30 minutes for a total incubation period of 5 hours.

### In vitro stripe assay

Protein carpets were generated using silicon matrices with a channel system as previously described (Knoll et al., 2007). The carpets consisted of alternating stripe patterns, where the first stripe contained a mixture of ephrin-Fc (or other ligand proteins) and Fc-specific Cy3 conjugate in a 4:1 weight ratio, and the second stripe contained only Fc reagents without the Fc-specific Cy3 conjugate. The e5 chick spinal motor column was dissected and collected as described in earlier studies (Gallarda et al., 2008; Kao and Kania, 2011). In brief, e5 chick embryos of either sex were collected in ice-cold motor neuron medium [Neurobasal (Invitrogen), B-27 supplement (1:50, GIBCO), 0.5 mM L-Glutamate (Sigma), 25 mM L-Glutamine (GIBCO), and Penicillin-Streptomycin (1:100, Wisent)]. The lumbar spinal cord was opened at the dorsal midline, allowing the excision of motor columns under a stereomicroscope. The spinal motor columns were identifiable as the bulging part of the open-book spinal cord. Sharp tungsten needles (World Precision Instruments) were used to remove the dorsal spinal cord at the lateral region, along with the floor plate and medial motor column (MMC) in the central region of the open-book spinal cord. The excised motor columns were trimmed into square-shaped explants, approximately one-fourth the width of the motor column. Between 10-20 explants were plated on laminin-coated (20 µg/mL, Invitrogen) 60-mm tissue culture dishes (Sarstedt), containing various combinations of protein stripe carpets in motor neuron medium. The cultures were incubated at 37°C in an atmosphere of 95% air and 5% CO_2_ for 18 hours.

### In situ mRNA detection and immunostaining

Chick and mouse embryos were fixed in 4% paraformaldehyde (Sigma) in PBS, equilibrated in 30% sucrose in PBS, embedded in O.C.T. (Sakura Finetek), and stored at −80°C. Sections of 12 micrometers were prepared using a cryostat microtome (Leica).

*In situ* mRNA detection was performed as previously described (Schaeren-Wiemers and Gerfin-Moser, 1993; Kania and Jessell, 2003). Briefly, tissue sections were first fixed in 4% paraformaldehyde in PBS for 10 minutes at room temperature, washed three times with PBS, and then digested in Proteinase K solution [1 µg/mL Proteinase K (Roche) in 6.25 mM EDTA (pH 8.0) and 50 mM Tris (pH 7.5) (Invitrogen)]. The sections were acetylated for 10 minutes by immersion in a solution containing 6 ml triethanolamine (Sigma), 500 ml double-distilled H**_2_**O, and 1.30 ml acetic anhydride (Sigma). After PBS washes, the sections were incubated in hybridization buffer [50% formamide, 5X SSC (0.75 M NaCl, 0.075 M NaAc), 5X Denhardt’s (Sigma), and 500 µg/mL Salmon sperm DNA (Roche)] for 2 hours at room temperature, followed by overnight incubation at 72°C with DIG-labeled RNA probes (prepared as described) at a concentration of 2-5 ng/µL in hybridization buffer. After hybridization, the samples were immersed in 5X SSC at 72°C, followed by two washes in 0.2X SSC at 72°C for 45 minutes each, and finally a wash in 0.2X SSC at room temperature for 5 minutes. The tissues were then rinsed with B1 buffer (0.1 M Tris, pH 7.5, and 0.15 M NaCl) for 5 minutes, blocked in B2 buffer (10% heat-inactivated horse serum in B1) for 1 hour at room temperature, and incubated with anti-DIG antibody (1:5000 in B2, Roche) overnight at 4°C. Samples were rinsed with B1, equilibrated with B3 buffer (0.1 M Tris, pH 9.5, 0.1 M NaCl, 0.05 M MgCl2), and incubated with B4 buffer [100 mg/mL NBT, 50 mg/mL BCIP (Roche), and 400 mM Levamisole (Sigma) in B3] in the dark to detect bound anti-DIG antibodies. The reaction was stopped by immersion in water.

Immunostaining was performed as described in previous studies (Kao et al., 2009; Kao and Kania, 2011). Briefly, sectioned tissues or cultured explants were washed in PBS and incubated in blocking buffer [1% heat-inactivated horse serum in 0.1% Triton-X/PBS (Sigma)] for 5 minutes, followed by overnight incubation at 4°C in the selected primary antibodies diluted in blocking solution. After primary antibody incubation, the samples were washed with PBS and incubated with appropriate secondary antibodies for 1 hour at room temperature. Refer to Table 1 for a list of the antibodies used.

**Table 1:**
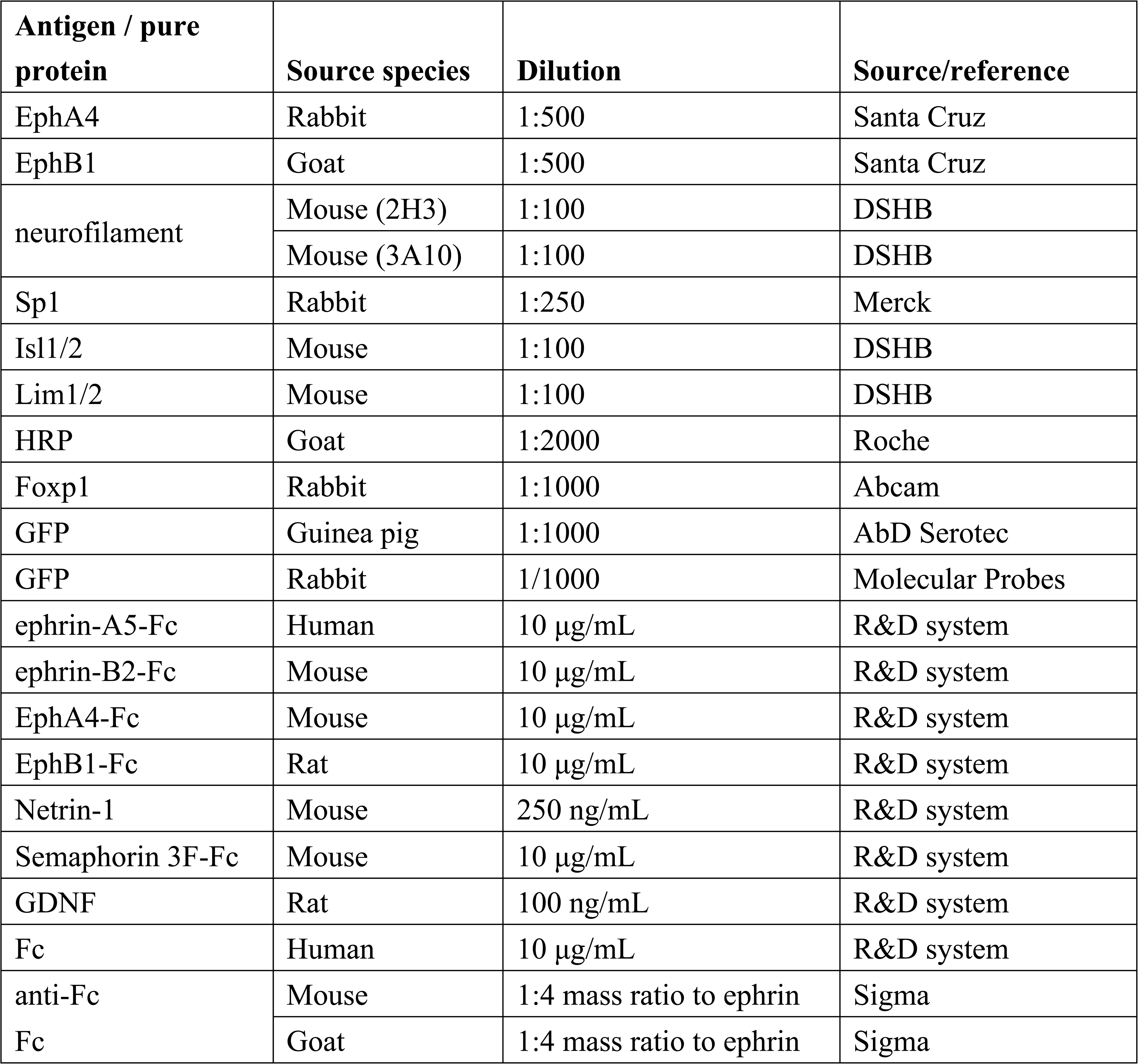
Antibodies and Fc reagents used.

| **Antigen / pure protein** | **Source species** | **Dilution** | **Source/reference** |
| --- | --- | --- | --- |
| EphA4 | Rabbit | 1:500 | Santa Cruz |
| EphB1 | Goat | 1:500 | Santa Cruz |
| neurofilament | Mouse (2H3) | 1:100 | DSHB |
|  | Mouse (3A10) | 1:100 | DSHB |
| Sp1 | Rabbit | 1:250 | Merck |
| Isl1/2 | Mouse | 1:100 | DSHB |
| Lim1/2 | Mouse | 1:100 | DSHB |
| HRP | Goat | 1:2000 | Roche |
| Foxp1 | Rabbit | 1:1000 | Abcam |
| GFP | Guinea pig | 1:1000 | AbD Serotec |
| GFP | Rabbit | 1/1000 | Molecular Probes |
| ephrin-A5-Fc | Human | 10 µg/mL | R&D system |
| ephrin-B2-Fc | Mouse | 10 µg/mL | R&D system |
| EphA4-Fc | Mouse | 10 µg/mL | R&D system |
| EphB1-Fc | Rat | 10 µg/mL | R&D system |
| Netrin-1 | Mouse | 250 ng/mL | R&D system |
| Semaphorin 3F-Fc | Mouse | 10 µg/mL | R&D system |
| GDNF | Rat | 100 ng/mL | R&D system |
| Fc | Human | 10 µg/mL | R&D system |
| anti-Fc<br>Fc | Mouse | 1:4 mass ratio to ephrin | Sigma |
|  | Goat | 1:4 mass ratio to ephrin | Sigma |

### Image quantification

Images were acquired using either a Leica DM6000 microscope or an EVOS FL microscope (Thermo Fisher). Quantification of GFP-labeled axonal projections was performed by thresholding and counting pixels in limb section images containing limb nerves (10-15 limb sections, each with a thickness of 12 micrometers), using Photoshop (Adobe). The dorsal or ventral limb nerve was identified by selecting the neurofilament channel and using the Lasso Tool, and pixel counts were obtained from the GFP channel by measuring the pixel intensity from the threshold to the maximal level, as indicated in the Histogram window. The number of motor neurons was quantified by counting cells in a series of spinal cord section images (5-10 sections from each embryo) using Photoshop. For quantification of GFP- or EphA4-labeled neurites in cultured motor neuron explants growing on different stripe types, over-threshold pixel counts were performed for the first and second stripe types across multiple images using Photoshop. To reduce potential experimental bias, multiple limb sections were selected at intervals of 2-3 sections along the anteroposterior axis to cover the entire crural plexus of limb nerves in each embryo. Similarly, cultured LMC explants were randomly selected for quantification. Additionally, the proportions of total and electroporated motor neurons were quantified in most experimental groups to ensure that there were no significant changes in cell identity or abnormal cell death, allowing for a reliable subsequent analysis of axon growth preferences (dorsal/ventral, medial/lateral, or first stripe/second stripe).

### Behavioral Experiments

For the behavioral assessments, including the open field test, Y-maze analysis, and grip strength test, the experimental mice were transported to the Taiwan Mouse Clinic at Academia Sinica, Taipei, Taiwan. The rota-rod test was conducted at the Animal Center of Taipei Medical University. An accelerating rota-rod (47650 Rota-Rod NG, Ugo Basile, VA, Italy) was used for this test, with the rotation speed gradually increasing from 4 to 40 rpm over the course of a 5-minute trial. Mice were placed on the rotating rod, and the time to fall was recorded. Each mouse underwent a training regimen consisting of three trials per day, with 30-minute intervals between trials, over the course of three consecutive days.

### Statistical analysis

Data from experimental replicate sets were analyzed using Microsoft Excel. The means of combined proportions or cell counts were compared using the Fisher’s exact test to evaluate the proportions of medial/lateral LMC neurons in retrograde labeling experiments, or the Mann-Whitney U test for comparing the proportions of dorsal/ventral preferences in limb nerves or cultured LMC neurites growing on 1^st^ stripe/2^nd^ stripe. The threshold for statistical significance was set at p < 0.05 (Poliak et al., 2015). For behavioral experiments, a two-sample paired t-test was used to compare the means of the data (Kao et al., 2015).

## Acknowledgement

This work was supported by the National Science and Technology Council, Taiwan (110-232-B-038-017-MY3), and the TMU Research Center of Cancer Translational Medicine from the Featured Areas Research Center Program within the framework of the Higher Education Sprout Project by the Ministry of Education in Taiwan. We thank Stephanie Ho for secretarial assistance. We thank the Taiwan Mouse Clinic, Academia Sinica and Taiwan Animal Consortium for the technical support in HCS, open field test, Y-maze analysis, and grip strength test. We appreciate that NLAC kindly provided the strain of C57BL/6-Sp1 tm1(GEMMS)Narl mice.

